# A guide to using a multiple-matrix animal model to disentangle genetic and nongenetic causes of phenotypic variance

**DOI:** 10.1101/318451

**Authors:** Caroline E. Thomson, Isabel S. Winney, Oceane C. Salles, Benoit Pujol

## Abstract

Non-genetic influences on phenotypic traits can affect our interpretation of genetic variance and the evolutionary potential of populations to respond to selection, with consequences for our ability to predict the outcomes of selection. Long-term population surveys and experiments have shown that quantitative genetic estimates are influenced by nongenetic effects, including shared environmental effects, epigenetic effects, and social interactions. Recent developments to the “animal model” of quantitative genetics can now allow us to calculate precise individual-based measures of non-genetic phenotypic variance. These models can be applied to a much broader range of contexts and data types than used previously, with the potential to greatly expand our understanding of nongenetic effects on evolutionary potential. Here, we provide the first practical guide for researchers interested in distinguishing between genetic and nongenetic causes of phenotypic variation in the animal model. The methods use matrices describing individual similarity in nongenetic effects, analogous to the additive genetic relatedness matrix. In a simulation of various phenotypic traits, accounting for environmental, epigenetic, or cultural resemblance between individuals reduced estimates of additive genetic variance, changing the interpretation of evolutionary potential. These variances were estimable for both direct and parental nongenetic variances. Our tutorial outlines an easy way to account for these effects in both wild and experimental populations. These models have the potential to add to our understanding of the effects of genetic and nongenetic effects on evolutionary potential. This should be of interest both to those studying heritability, and those who wish to understand nongenetic variance.

## Introduction

Genetic and nongenetic sources of phenotypic variation and similarity between individuals are often studied in isolation, with different research aims. Laboratory experiments have demonstrated genetic and nongenetic sources of similarity (e.g. Cubas *et al.* 1999), but the use of fixed genetic and/or environmental backgrounds (e.g. epigenetic recombinant inbred lines; Latzel *et al.* 2012; Cortijo *et al.* 2014) can limit their relevance in nature. Studies in wild populations are more biologically realistic, but cannot control for all sources of variation. For example, such studies may investigate quantitative genetic (Falconer & Mackay 1996; Lynch & Walsh 1998), social (Otto *et al.* 1994; Grant & Grant 1996), or environmental sources of variation (Legendre 1993). Yet, these mechanisms are likely to act simultaneously on phenotypes, which makes them challenging to disentangle. Whilst there are attempts to consider genetic and nongenetic aspects of phenotypic variation simultaneously (e.g. Day & Bonduriansky 2011; Townley & Ezard 2013), these methods are challenging to apply with current empirical data. Thus there is a need to distinguish multiple sources of variation within a single model framework that is simple to use with realistic data. This will be valuable for addressing questions about how phenotypes are shaped, both in wild and experimental settings, and the evolutionary consequences of these causes of variance.

### Quantitative genetic components of phenotypic variance

Quantitative genetics assesses the extent to which related individuals show similar phenotypic traits, making the assumption that this is because of shared genetic effects (Falconer & Mackay 1996). Furthermore, this assumes that continuous phenotypic traits are determined by a large number of genes with small additive effects (the infinitesimal model; Falconer & Mackay 1996; Lynch & Walsh 1998), and environmental influences. The ‘animal model’ is a powerful tool in quantitative genetic, used to disentangle the origins of phenotypic variation in natural systems (Kruuk 2004). Adopted from animal breeding studies, this mixed-effects model associates variation in phenotypic traits with relatedness information to give an estimate of the (additive) genetic variation underlying the trait (Wilson *et al.* 2010). From this, researchers primarily calculate narrow-sense heritability, the portion of population phenotypic variance attributable to additive genetic variation, which provides predictions about a population’s expected response to selection (Lush 1937; Lande 1979). Therefore, the animal model provides a link between phenotypic traits and their evolutionary potential.

The animal model is mostly used to estimate additive genetic variance (and heritability), but phenotypic variance can sometimes be affected by other factors that reduce the precision of these estimates. Genetic variance can include dominance and epistatic variance (Falconer & Mackay 1996), in addition to the additive genetic variance, meaning that additive genetic variance may be overestimated. Dominance variance can be partitioned by comparing full- and half-siblings, or by including a dominance matrix in the animal model, although this is rarely done in wild studies (Wolak & Keller 2014).

Nongenetic factors can also cause misestimation of additive genetic variance in some systems. The main assumptions underlying the animal model are that relatives share genes, and therefore the genetic effects that shape their genotype, but that any effects shaping unexplained variance are independent between individuals (Lynch & Walsh 1998). Shared genes are not the only sources of similarity between individuals, however, which violates these assumptions (Kruuk & Hadfield 2007). These sources of variation may need to be accounted for to accurately estimate heritability and evolutionary potential, and can be of interest in their own right. In this paper we focus principally on three nongenetic sources of variation: environmental, epigenetic, and social sources of variation, although others are known to exist (Danchin *et al.* 2011). Whilst environmental sources of variation, including the environment caused by social partners, have previously been included in the animal model in various ways, variation attributable to epigenetics or social networks have not. We begin, therefore, by briefly reviewing how environmental sources of variance can be included in quantitative genetic models.

### Including the environment in the animal model

Shared environments can increase the phenotypic resemblance between relatives and alter heritability estimates. Shared rearing environments in particular can increase sibling similarity. To address this problem, the common environmental factor causing individuals to resemble one another can be included as a random effect in the animal model. For example, models may include the year of measurement (e.g. Reale *et al.* 2003) or year of birth (e.g. Charmantier *et al.* 2006) to account for broad-scale environmental variance. Shared rearing effects can also be accounted for in this way (e.g. Kruuk *et al.* 2001; Merilä & Sheldon 2001), particularly when combined with cross-fostering experiments (Kruuk & Hadfield 2007; Thomson *et al.* 2017).

In many cases, some portion of the environment that an individual experiences can be attributed to other individuals. For example, shared rearing environments can include environmental parameters generated by one or both parents (Mousseau & Fox 1998; Green 2008). Parental effects can be accounted for in simple cases by including the identity of one or both parents (usually the mother) as an additional random effect (e.g. McAdam *et al.* 2002; Noble *et al.* 2014; McFarlane *et al.* 2015). This accounts for trait variance that can be attributed to the parent(s) (parental performance; Willham 1963, 1972), rather than attempting to identify and account for all parental phenotypic traits that cause the focal trait variance (although see Noble *et al.* (2014)).

More generally, the variance in a focal individual’s phenotype can be affected by indirect genetic effects (IGEs; Griffing 1976; Moore *et al.* 1997; Wolf *et al.* 1998). These arise when a focal individual’s environment is altered by a conspecific (whether parental or not), as determined by the conspecific’s own genes. When IGEs occur, the residual variance in a trait is partially heritable and can affect evolutionary trajectories (Bijma 2014). Animal models that incorporate IGEs have been developed (Muir & Schinckel 2002; Arango *et al.* 2005; Muir 2005; Van Vleck & Cassady 2005; Bijma *et al.* 2007), and implemented within the fields of animal and plant breeding, allowing better estimation of the total heritabilities of traits. Additionally, experimental populations have been used to estimate IGEs on aggression (Wilson *et al.* 2009; Santostefano *et al.* 2016). In wild populations, there have been some estimates of IGEs, particularly where only one or two individuals have indirect genetic effects on traits. Parental effects can be estimated as parental genetic effects, wherein the covariance between the effects of mothers (or fathers) is determined by the additive genetic relatedness matrix, analagous to the estimation of (direct) additive genetic variance (Mrode & Thompson (1996); Lynch & Walsh (1998); Kruuk (2004); for examples see: Wilson *et al.* (2005a); McFarlane *et al.* (2015)). Additionally, wild studies have estimated IGEs of males on female reproductive timing (Brommer & Rattiste 2008; Liedvogel *et al.* 2012), and those of helpers at the nest in long-tailed tits (*Aegithalos caudatus*; Adams *et al.* 2015).

### Accounting for spatial similarity in environments

Environments are often heterogeneous over both space and time (e.g. Koenig 1999; Dormann *et al.* 2007), and may show spatial (and temporal) autocorrelation. Factorial random effects such as birth year may not capture the variance due to continuous and spatially autocorrelated environmental effects. This can lead to the misestimation of variance components, particularly if relatives share environments due to limited dispersal. Within quantitative genetics, spatial autocorrelation of phenotypes can be accounted for in the animal model. Autoregressive (AR) models account for the similarity between individuals that are close in space, using a two dimensional row-by-column model, first incorporated into animal models for agricultural trials (Gleeson & Cullis 1987; Cullis & Gleeson 1991. This method has been widely used in agriculture, including in forest genetic trials (Magnussen 1993; Dutkowski *et al.* 2002), although the AR term has not always accounted for all environmental variance (Dutkowski *et al.* 2006). In one recent paper, Costa e Silva *et al.* (2017) included both IGEs of nearest neighbours and spatial effects in models of leaf disease and growth in *Eucalyptus* to show that these processes may act simultaneously and interact.

The effect of spatial autocorrelation on heritabilities in the wild has received relatively little attention, although some studies examining its effects are emerging. Van Der Jeugd & McCleery (2002) used parent-offspring correlations to show that including spatial autocorrelation reduced estimates of laying date heritability in great tits (*Parus major*). Subsequently, three studies have included spatial autocorrelation in animal models for wild populations (red deer, *Cervus elaphus*; Stopher *et al.* 2012; song sparrows, *Melospiza melodia*; Germain *et al.* 2016; Soay sheep, *Ovis aries*; Regan *et al.* 2017). Whilst Stopher *et al.* (2012) found relatively large variance components associated with spatial autocorrelation, particularly for home range size traits, such large variance components were not found by Regan *et al.* (2017) for morphological or phenological traits, or by Germain *et al.* (2016) for breeding time.

### An alternative approach to measuring environmental similarity

Three of the papers above (Stopher *et al.* 2012; Germain *et al.* 2016; Regan *et al.* 2017) demonstrate an additional way to account for variance caused by the environment, which we build on in this paper. This technique was first applied by Stopher *et al.* (2012), who included a matrix of home-range overlap (the “S” matrix) in animal models. Spatial observations were used to estimate an individual’s home range, and the overlap of individual’s home ranges were estimated and used to generate the S matrix. The S matrix was then incorporated into the animal model to estimate trait variance due to shared space. This model had the expectation that if the environment affected trait values, individuals that shared space would be similar in their trait values. The S matrix would therefore account for the variance associated with environmental effects. The variance accounted for by the S matrix was generally comparable to the magnitude of spatial autocorrelation in both red deer (Stopher *et al.* 2012) and Soay sheep (Regan *et al.* 2017). Germain *et al.* (2016) used a similar method to estimate shared environmental effects on breeding date in song sparrows, using overlap in buffer zones around nest boxes. As with spatial autocorrelation, there was no significant effect of nest area overlap on breeding date heritability estimates.

### Broader applications of multiple matrices

The S matrix will capture environmental sources of trait variance when it appropriately captures environmental heterogeneity. However, individuals that do not share space may still experience similar environments, whereas those that do share space may differ in the environments that they experience. We propose that similar matrices could be built using multiple measures of the environment itself, to describe the similarity in environment that individuals experience, and account for environmental variance in the animal model. The formation and use of such a matrix is demonstrated in the tutorial section below and in the supplement.

We show two additional variance parameters that can be estimated using these matrices, beyond environmental variance. Firstly, phenotypic variation can also be caused by epigenetic variation (Jablonka & Raz 2009; Lira-Medeiros *et al.* 2010). This can be induced by environmental conditions (Foust *et al.* 2016), but has also been found in the absence of genotypic and environmental variation (Wong *et al.* 2005; Peaston & Whitelaw 2006). Estimates of additive genetic variation are made from statistical patterns that can include additive epigenetic variation, particularly if epigenetic marks are inherited (for which there is experimental evidence; Champagne & Meaney 2006; Paszkowski & Grossniklaus 2011). To separate additive genetic and epigenetic variation, inclusion of matrices describing epigenetic “relatedness”, or similarity, between individuals should allow estimation of epigenetic variance.

Secondly, the social environment can affect phenotypic traits. An individual can learn behaviours from others with whom it interacts (e.g. Waal *et al.* 2013; Aplin *et al.* 2015). When individuals are embedded within a social network, information can be transmitted more strongly amongst frequently interacting individuals (Franz & Nunn 2009; Allen *et al.* 2013). If relatives are social partners (e.g. Hirsch *et al.* 2012; Holekamp *et al.* 2012), this can potentially inflate estimates of genetic variance in an animal model. Conversely, to understand social effects on trait values, researchers should take relatedness and additive genetic variation into account. We propose using a matrix to account for links between individuals in a social network, and thereby estimate the variance associated with social structures in an animal model. Inclusion of social information into an animal model has previously been attempted for bottlenose dolphins (*Tursiops* sp.; Frère *et al.* 2010), although this study had methodological issues, and inclusion of both genetic and social information within a single model was not attempted.

The purpose of this paper is to outline the tools available to disentangle causes of genetic and nongenetic resemblance between individuals, within the animal model framework. We take advantage of the versatility of this model to demonstrate the use of the multiple matrix method proposed by Danchin *et al.* (2011), and first implemented for nongenetic data by Stopher *et al.* (2012). Matrices that account for nongenetic causes of similarity between individuals are included in the same way as a matrix of genetic relatedness, and can be derived from multiple sources of similarity, including those described above. Using simulated data we provide a complete tutorial, with code provided in the supplementary materials, to demonstrate how to carry out such analyses with R packages (R Core Team 2016). We show how phenotypic variance can be partitioned into genetic and nongenetic components, including parental effect variance, in three different phenotypic traits. We also give a qualitative overview of the data sizes needed to run these models. This extension of the animal model has the potential to bridge multiple areas of research on genetic and nongenetic sources of phenotypic variation.

## Tutorial

The animal model is a mixed-effects model (i.e. a linear regression containing fixed and random terms), that partitions the total phenotypic trait variance in a population (*V^p^*) between various sources, including additive genetic variation (*V_a_*). The animal model is a powerful tool for disentangling multiple sources of variation, both genetic and nongenetic. For readers who are new to the animal model, we recommend Lynch & Walsh (1998), Kruuk (2004), and Wilson *et al.* (2010) as an introduction.

Our aim here if to demonstrate the creation and use of nongenetic matrices in animal models. To illustrate this, we simulated three phenotypic traits, and ran animal models on them, as detailed below. Animal models were run using the R package ASReml-R (Butler 2009). Full details of the simulations are given in the Supplementary Information, with a brief overview in the section below. The subsequent sections contain the theory behind the variance components included, followed by the results of models that used matrices of environmental similarity. We show how matrices derived from environmental, epigenetic and social network information can be included to estimate variance. A step-by-step guide with code (for ASReml-R) is given in the Supplementary Information, with additional information on how to carry out these analyses in MCMCglmm (Hadfield 2010), a Bayesian R package. Fitting some or all of these models may be possible in other software and R packages, including standalone ASReml (Gilmour *et al.* 2009), Wombat (Meyer 2007) and INLA (Holand & Martino 2016). Finally, to provide an idea of the data requirements of these models, we show the effects of data removal on the model in the Supplementary Information.

### Data Simulation

To demonstrate how to use multiple-matrix animal models, we provide a simulated dataset of a population of merpeople (mermaids and mermen). This imaginary population contains ten semelparous generations, of between 150 and 250 individuals per generation (total population size 1806). Half of each generation were selected and assigned to breeding pairs at random (the other half were non-breeding), and offspring were then randomly assigned to these pairs. To simulate extra-pair mating, 10% of offspring had one parent reselected from the breeding population.

To simulate environments and their effects, each individual was assigned a location on a 50×50 grid, with the first generation positioned on the grid at random. Subsequent generations dispersed from their mothers, following a lognormal distribution function (*μ* = 1, *σ*^2^ = 1, see supplementary information). Five continuous environmental variables (temperature, salinity, pH, coral coverage, depth) were simulated, with spatial autocorrelation across the 50×50 grid (the covariance between locations was determined by the matrix **e**^−**ϕD**^, where **D** is a matrix of distances and *ϕ* = 0.15). The five environmental variables were independently simulated and had negligible covariances. Individuals were then assigned a set of environmental variables according to their location on the grid.

We also simulated a set of epialleles, to show how an epigenetic similarity matrix can be used in an animal model. We simulated 230 CpG islands (regions of the genome with high G and C content; Gardiner-Garden & Frommer 1987), wherein the number of methylated CpG sites within each island was allowed to vary between 0 and 20. For 100 of these, the methylation was affected by the environmental variables an individual experienced itself. For 30, methylation was affected by either the maternal environment, or was reset to the value determined by the individual’s own environment (with 50% probability of reset), such that epigenetic marks could be transmitted across generations. The final 100 islands were uninherited and unaffected by the environment, so each individual had a randomly generated value between 0 and 20 for each island.

To demonstrate how social effect variance can be accounted for, we simulated a social network of within- and between-generation interactions, based on the spatial locations of individuals. Within generations, the probability of connection in the social network declined with distance. Between adjacent generations, there was a weaker probability of connections, although the probability remained higher for mother-offspring pairs. Non-adjacent generations had no direct links in the social network.

#### Trait values

We simulated three continuous phenotypic traits, composed of different sources of variance. Each individual variance component was simulated with a mean of zero and variance of 1. The total variance of each trait therefore depended on the number of components used to generate it. All trait values included additive genetic and residual effects. Each environmental variable (*k_l_*, where *l* indexes environments) had a linear effect on the trait (*β_l_*), such that the total effect of the environment on the trait for individual *i* is 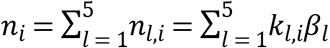
, with variance 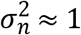. Direct effects of epigenetic marks or the social network were not simulated, but were expected to account for environmental variance (Fig 1). No covariances were simulated between variance components.

**Fig 1:**
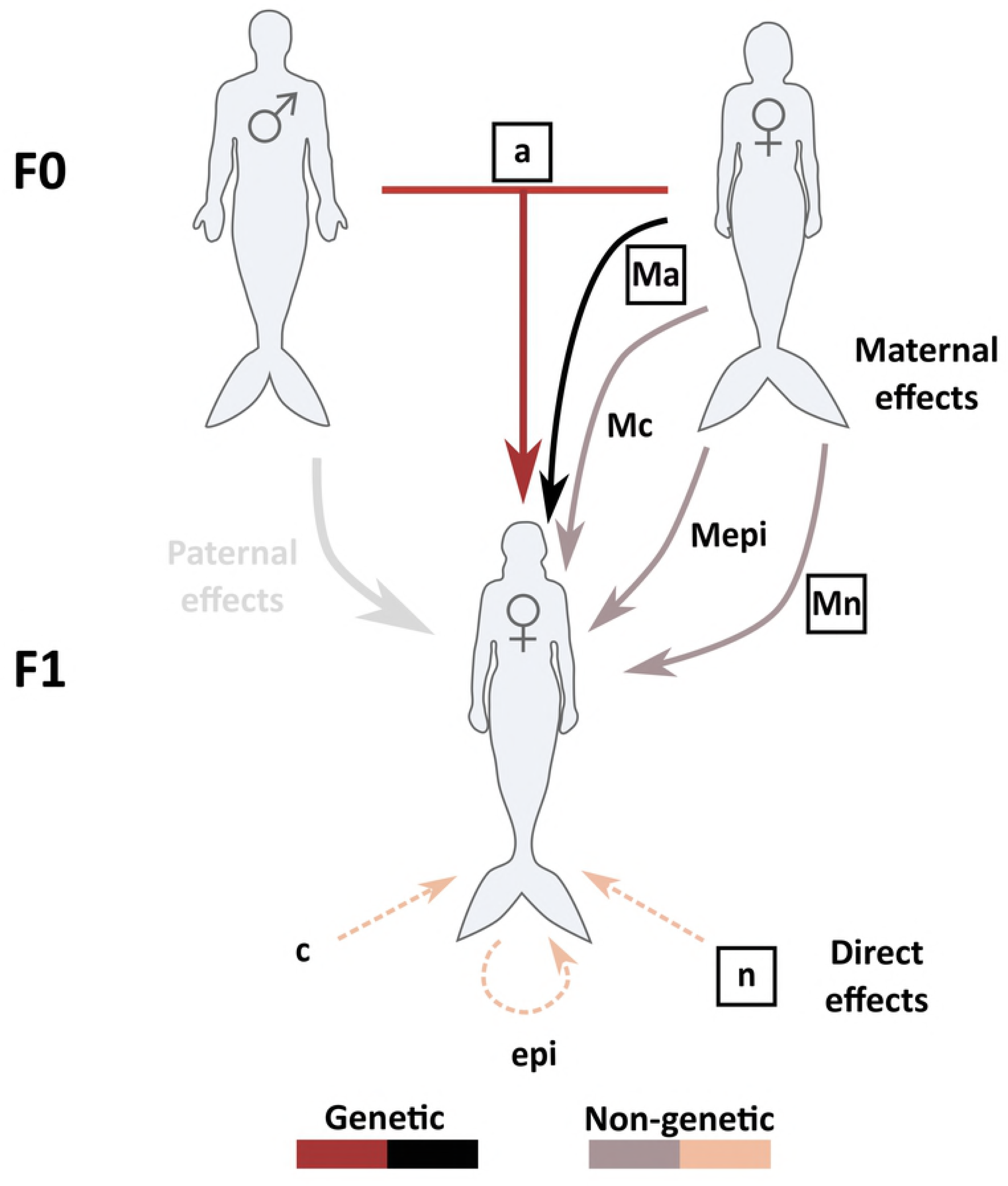
The potential inputs that could affect trait values in phenotypic traits in merpeople. Boxed parameters are the true parameters included in at least one of the traits. Unboxed parameters are included in models as potential sources of variance, but did not affect the simulated traits. *a* represents additive genetic effects, and *Ma* the maternal genetic effect. *n* is the total direct environmental effect, and *Mn* the maternal environmental effect. *epi* and *Mepi* are putative direct and maternal epigenetic effects, and *c* and *Mc* are putative direct and maternal social network/cultural effects. Paternal effects were not included in the simulations or models, but are shown here as a potential effect on trait values. F0 and F1 indicate generations.

The first trait was tail-fin colour. We use this trait to illustrate how the additive genetic and environmental variance can be confounded in an animal model, and how environmental variance can be accounted for. Individual trait values for tail-fin colour (*y_i_*_,1_) were:

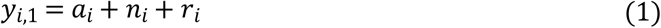

where *a_i_* is the additive genetic effect, *n_i_* is the effect of the five environments, and *r_i_* is a residual effect. Trail-fin colour has a raw trait variance 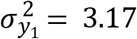
, and an expected heritability of ~0.33.

Trait two was body size (*y*_2_). This trait is used to illustrate maternal variance, particularly how it can be estimated with an environmental matrix. No paternal effects were simulated or modelled. Trait values were:.

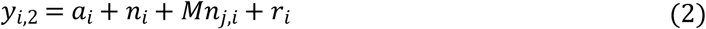

where maternal environmental variance 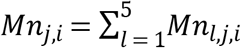
 (where *j* indicates the mother of individual *i*). The effect a mother had on her offspring’s trait was determined by the environment she experienced, akin to direct environmental effects. Total raw trait variance was 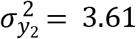
, with an expected heritability of ~0.25.

The third trait, swimming speed (*y*_3_), was used to illustrate the partitioning of different sources of maternal variance. This trait had all the components above, and also contained maternal genetic variance (*Ma_j_*_,*i*_), wherein the effect of a mother was also affected by her own genes:

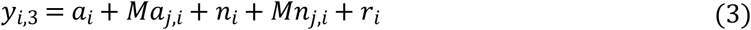

Total raw variance was 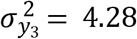
, with an expected heritability of ~0.2.

### Tail-fin colour - estimating additive genetic and environmental variance

#### Additive genetic variance

In a basic animal model, an individual (*i*) has a trait value (*y_i_*) that is is composed of at least the three following elements: the population mean (*μ*), the breeding value of the individual (*a_i_*, or the effect of the genotype relative to the population mean) and a residual (*r_i_*), such that the total trait is *y_i_* = *μ* + *a_i_* + *r_i_*. In a more general form, this can be rewritten as:

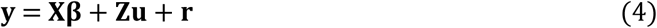

where **y** is a vector of trait values in all individuals, **X** is the design matrix relating fixed effects to individuals, and **β** is a vector of fixed effects. In the case the models here, which contains the intercept and no fixed effects, **X** is a vector of ones, and **β** =*μ*. **Z** is the design matrix relating random effects to each individual, and **u** is a vector of random effects.

When only one random effect (the additive genetic effect) is included, **Z** becomes the identity matrix (**I**) and **u** is a vector of additive genetic effects. Both **u** and and **r** are assumed to be normally distributed with means of zero, and variances of *V_a_* and *V_r_*, respectively (i.e. **u** ~ *N*(0,*V_a_*), **r** ~ *N* (0,*V_r_*)). Residual errors are assumed to be independent between individuals, so that the variance-covariance matrix for **r** is **R** = **I** \**V_r_*. Taking **G** to be the variance-covariance matrix for **u**, this can be derived from the expected covariance between relatives in additive genetic effects **G**= **A** * *V_a_***. A** is the additive genetic relatedness matrix (Fig 2a), which is derived from the pedigree or genomic relatedness data (Wilson *et al.* 2010; Bérénos *et al.* 2014). This matrix contains the elements *A_i_*_,*j*_ = 2*θ_i_*_,*j*_, where *θ_i_*_,*j*_ is the coefficient of coancestry (the probability that an allele in individual *i* is identical by descent to one of the alleles at the same locus in individual *j*). Thus for any pair of individuals, the additive genetic covariance between them is *G_i_*_,*j*_ = 2*θ_i_*_,*j*_\**V_a_*.

**Fig 2:**
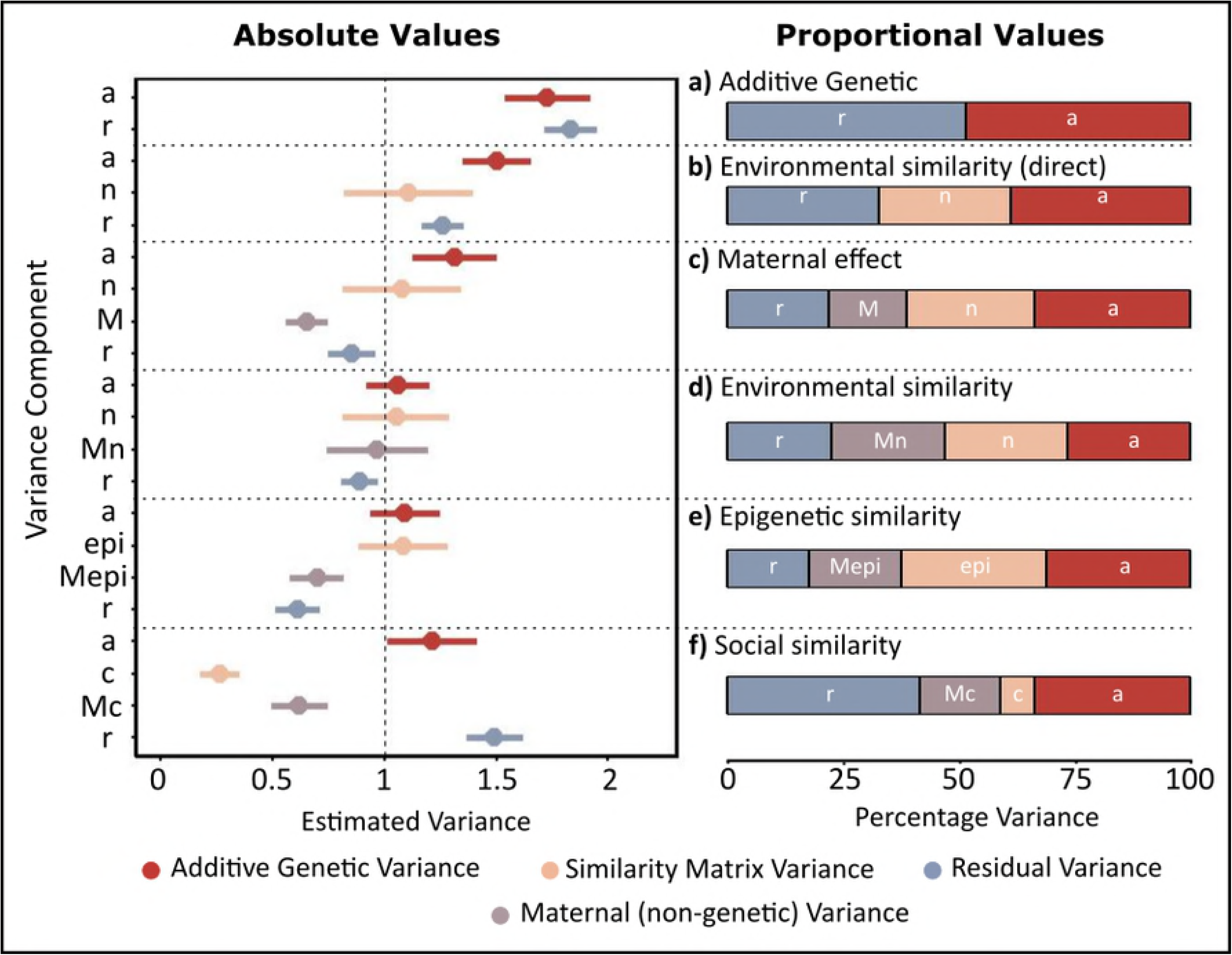
Reduced versions of the matrix information included in the animal models shown in this paper (a) The genetic relatedness matrix, derived from a pedigree, which contains the probabilities that individuals *i* and *j* share alleles that are identical by descent. (b) The environmental similarity matrix, which contains elements *s_n_*_(*i*,*j*)_ (written in fig 2 as *S*, as the same equation is used for elements of *S_epi_*) measuring the Euclidean distance between the environment individuals experience, *p* and *q* being the scaled environmental vectors for individuals *i* and *j* across *l* environments. (c) Epiallelic similarity, which contains elements *S_epi_*_(*i*,*j*)_, measuring the Euclidean distance between individuals’ epigenetic profiles, where *p* and *q* are the scaled epigenetic vectors for individuals *i* and *j*, across *l* epigenetic islands. (d) The geodesic distance matrix (social network connectedness) that shows the proximity of individuals in the social network, based on minimum path lengths (*g_i_*_,*j*_).

Using the additive genetic relatedness matrix, the animal model relates trait covariance between individuals to their relatedness, and thus we are able to separate the additive genetic variance (*V_a_*, variance in *a_i_*) contributing to a trait from the residual variance (*V_r_*, variance in *r_i_*). The phenotypic variance (*V_p_*) is then the sum of the additive genetic and residual variances:

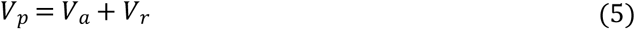

#### Environmental variance

Nongenetic factors can also cause individuals to resemble one another. To account for this, we can include information on nongenetic similarity between individuals in the model, with a method analogous to the use of additive genetic relatedness.

Shared environments can induce similar trait values between individuals, and thus create patterns of phenotypic variation that we may want to estimate. If related individuals share the same environment, neglecting shared environmental effects can inflate the estimated additive genetic variance (*V_a_*). Environmental causes of resemblance can be included as fixed or random effects in the model (Kruuk & Hadfield 2007), such as a discrete environment classification (nest box, year of birth etc.). In this paper, however, we are more interested in how multiple continuous environmental parameters could be used to account for phenotypic variation. We do this by including a matrix that describes environmental similarity between individuals. To separate the shared environment variance from additive genetic variance in the model, the data requires that there are related individuals found across different environments (Danchin *et al.* 2013). However, if relatives all live in different environments, environmental effects will be difficult to distinguish from residual variation. Thus, these models also require that several group members are found in the same environment (Kruuk & Hadfield 2007).

We took a measure of the similarity of the environment shared by each pair of individuals (across all generations), which generates the variance-covariance matrix associated with an environmental random effect that can be included in the animal model. We call this the **S_n_** matrix, sensu Stopher *et al.* (2012), with subscript *n* indicating environmental effects. Similarity was calculated from the five continuous environmental measures that were simulated. These environmental measures were first centered and scaled, so that all had a mean of zero and variance of one. For each pair of individuals, the Euclidean distance between environmental measures was then calculated. This is equal to the straight-line distance between the two vectors of environmental measures in multivariate space, and so describes the similarity in environmental measures between individuals 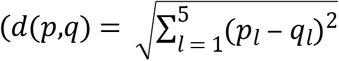, where *p* and *q* are the vectors of centered and scaled environmental measures for individuals *i* and *j*). This was scaled so that the **S_n_** matrix contains values between 0 (dissimilar environments) and 1 (identical environments), with 1’s on the diagonal because individuals have an identical environment to themselves (Fig 2b). Thus, each element of the matrix is:

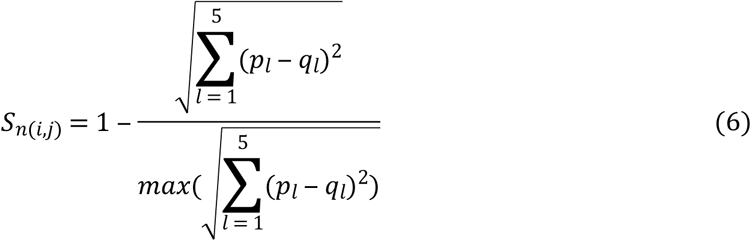

Inclusion of the **S_n_** matrix as the variance-covariance matrix for an environmental random effect relates shared environments with phenotypic traits, and evaluates whether individuals experiencing similar environments have similar trait values. Environmental effects (*n*) are assumed to be normally distributed with mean of zero, **n ~** *N*(0,**S_n_***V_n_*). This gives:

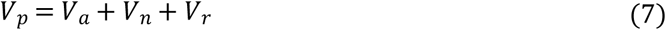

#### Variance estimates for tail-fin colour

Applying a basic animal model of the form in equation 5 for tail-fin colour gave a significant estimate of additive genetic variance that was larger than the simulated variance of 1 (*V_a_* = 1.714, SE=0.19, p<0.001). Similarly, the residual variance was larger than the simulated value of 1 (*V_r_* = 1.714, SE=0.11). The model therefore gives an overestimate of trait heritability *h*^2^ = 0.538 (SE=0.042; Fig 3a). This overestimation is due to the fact that there were also direct environmental effects on the trait value that were not accounted for.

The environmental similarity matrix was subsequently included in the animal model for tail-fin colour, and produced accurate variance estimates (Fig 3b). As this trait was composed of additive genetic and environmental effect, this represents the complete (and correct) model of phenotypic variance. All three components were simulated to have variances around 1, and the heritability around 0.33. The values estimated in this model were close to these values, confirming the efficacy of this method; *V_a_* = 1.1 (SE=0.12, p<0.001), *V_n_* = 0.986 (SE=0.22, p<0.001), *V_r_* = 0.894 (SE=0.07), with *h*^2^ = 0.369 (SE=0.042).

**Fig 3:**
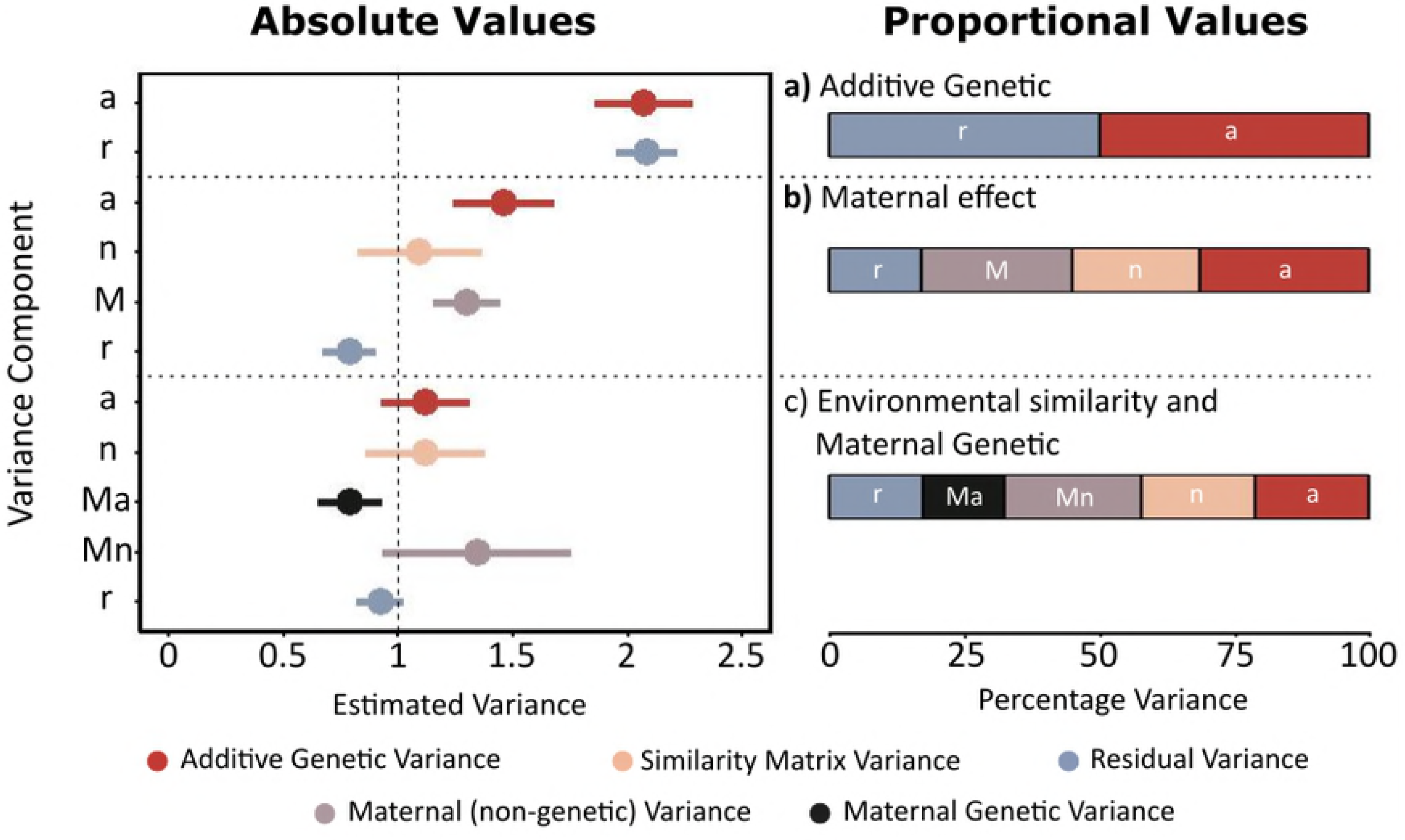
Variance components of tail-fin colour (trait 1), which was simulated to contain additive genetic, environmental, and residual variance, each with mean=0 and variance=1. For each of the models, the pair of plots show the absolute values (left plot) and the proportion (right plot) of total trait variance explained. a) Model containing only additive genetic (*a*) and residual variance (*r*). b) Model including additive genetic (*a*), environmental (*n*), and residual variance (*r*) (the true model). c) Model including additive genetic (*a*), epigenetic (*epi*), and residual variance (*r*). d) Model containing additive genetic (*a*), social (*c*), and residual variance (*r*). The vertical dashed line shows simulated value of one for all variance components.

### Body size - estimating maternal environmental variance

#### Including Parental Effects

Parents can form a portion of the environmental causes of trait values (i.e. an external input to the focal trait). Thus, we can think of parental effects as the combined influence of one or more (unmeasured) parental traits that affect the offspring’s phenotype (e.g. feeding rate), influenced by the parents own genotypes and/or environments. Parental effects can inflate the similarity between relatives, as siblings generally experience the same parental effects.

We can account for parental effects in the animal model. In the simplest case, this is done by including parental identity as an additional random effect. This requires that the parents have more than one offspring in the data, and that half-siblings are also present. Maternal half-siblings share both additive genetic and maternal effects, whereas paternal half-siblings share additive genetic and paternal effects. Information also comes from individuals that are related but with different mothers. We show maternal effects here, but the equivalent can be done for paternal effects. With the inclusion of maternal variance (*V_M_*), the total variance estimated in the model becomes:

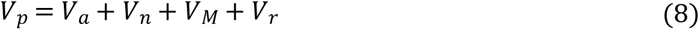

This model assumes that there is no covariance between mothers in the effects that they have on their offspring. However, mothers have not been simulated to be independent in their effects on offspring body size. This arises due to shared environments - mothers with similar environmental values are expected to express similar maternal effects on their offspring. For example, mothers that share environments may access similar types and quantities of food, and therefore show similar provisioning. We can account for this non-independence by estimating maternal environmental variance (*V_Mn_*), using the same matrix of environmental similarity (*S_n_*) as was used for the direct environmental effect. The total phenotypic variance is then estimated as:

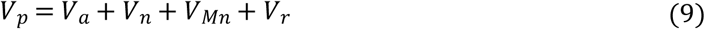

#### Variance estimates for body size

Body size was simulated to contain the same variance components as tail-fin colour, but with the addition of maternal environmental variance (*V_Mn_*; equation 2). In an animal model estimating only additive genetic and residual variances, both variances were overestimated, *V_a_* = 1.73 (SE=0.19, p<0.001), *V_r_* = 1.833 (SE=0.12; Fig 4a). Accordingly, the heritability estimate is also upwardly biased, *h*^2^ = 0.486 (SE=0.04).

**Fig 4:**
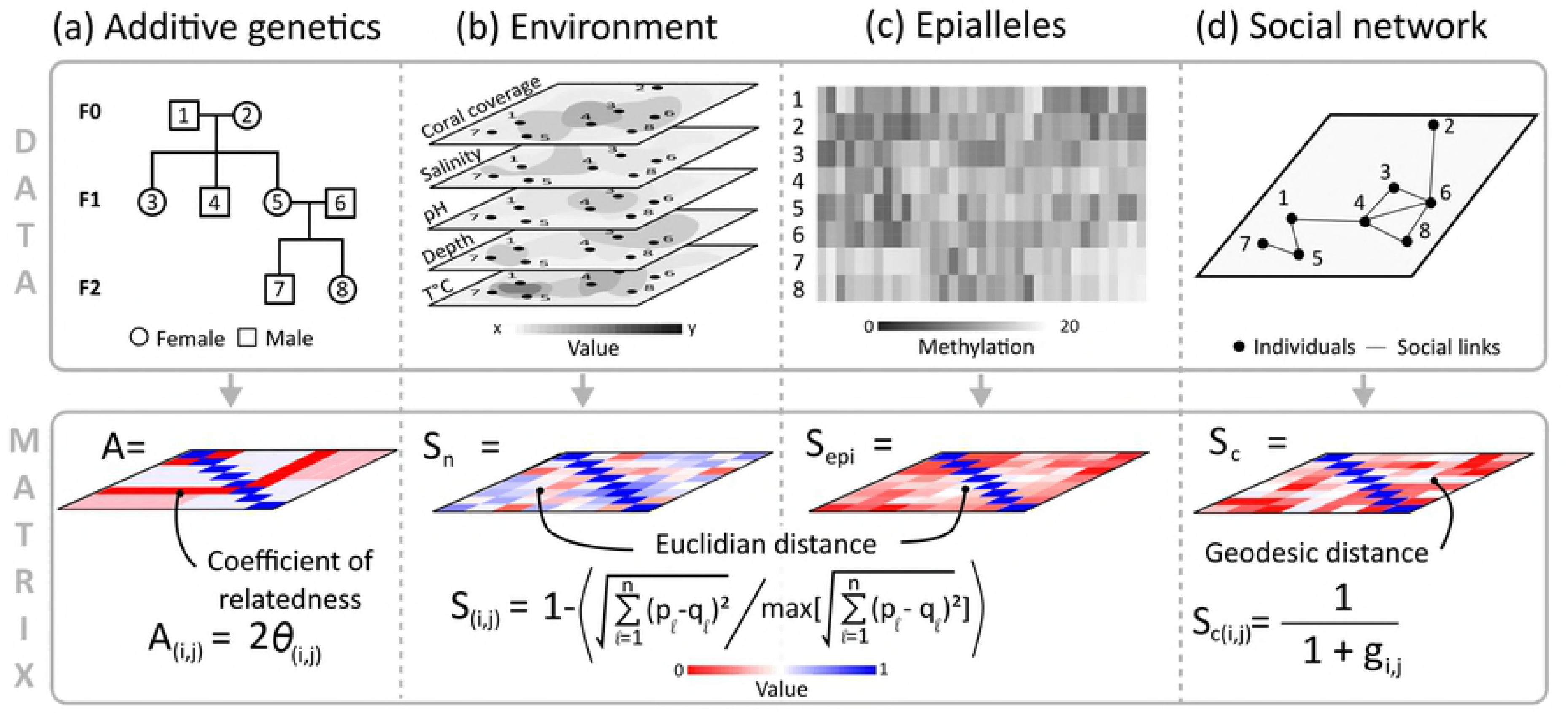
Variance components of trait 2. This trait was simulated to contain additive genetic, environmental, maternal environemental, and residual variance, each with mean=0 and variance=1. For each of the models, the pair of plots show the absolute values (left plot) and the proportion (right plot) of total trait variance explained. a) Model containing only additive genetic (*a*) and residual variance (*r*). b) Model including additive genetic (*a*), environmental (*n*), and residual variance (*r*). c) Model including additive genetic (*a*), environmental (*n*), and residual variance (*r*), and maternal variance estimated by including maternal identity as a random effect (*M*). d) Model including additive genetic (*a*), environmental (*n*), maternal environmental (*Mn*), and residual variance (*r*) (the true model). e) Model including additive genetic (*a*), epigenetic (*epi*), maternal epigenetic (*Mepi*), and residual variance (*r*). f) Model containing additive genetic (*a*), social (*c*), maternal social (*Mc*), and residual variance (*r*).

Including the environmental matrix in the animal model, we estimated a significant amount of (direct) environmental variance in the trait (Fig 4b), *V_n_* = 1.108 (SE=0.29, p<0.001). This somewhat reduces the estimate of additive genetic variance *V_a_* = 1.503 (SE=0.15, p<0.001), residual variance *V_r_* = 1.259 (SE=0.09) and heritability *h*^2^ = 0.388 (SE=0.041). Nevertheless, some variance estimates remain upwardly biased because maternal effects were not accounted for.

Including a simple maternal effect in the animal model (as in equation 8), we found significant maternal effect variance *V_M_* = 0.653 (SE=0.09, p<0.001), although it was underestimated relative to the true simulated variance. This reduced the estimate of the additive genetic (*V_a_* = 1.314, SE=0.19, p<0.001), and residual variance (*V_r_* = 0.855, SE=0.1), although these remain overestimated, as was heritability (*h*^2^ = 0.337, SE=0.046; Fig 4c). Environmental variance was unchanged compared to the previous model (*V_n_* = 1.079 SE=0.27, p<0.001).

The misestimation of the variance components in the model above was due to the non-independence of mothers in the effects they had on offspring body size. Therefore, we ran a model including maternal environmental variance (as in equation 9). We found a significant maternal environmental effect, *V_Mn_* = 0.97 (SE=0.23, p<0.001), similar to the simulated variance of one. Inclusion of the maternal environmental effect led to variance estimates close to 1 for all components (Fig 4d): *V_a_* = 1.059 (SE=0.14, p<0.001), *V_n_* = 1.052 (SE=0.24, p<0.001), and *V_r_* = 0.892 (SE=0.08). The heritability of the trait was *h*^2^= 0.267 (SE=0.038), similar to the expected heritability of 0.25. Thus, the model that matched the components of the simulated trait estimated the variance components.

### Swimming Speed - including maternal genetic variance

The final simulated trait, swim speed, is composed of the same components as body size, with the addition of maternal genetic effects (equation 3). We use this trait to demonstrate the difficulty of disentangling maternal genetic and environmental effects.

As with maternal environmental variance, maternal genetic variance causes mothers to covary in their maternal effects. To estimate maternal (or paternal) genetic effects using relatedness derived from a pedigree, the data needs to contain at least three generations of individuals (Wilson *et al.* 2010). To account for maternal genetic effects in the animal model an additional random effect is included, with a variance-covariance matrix determined by the additive genetic relatedness matrix, as is done for additive genetic variance (Mrode & Thompson 1996; Lynch & Walsh 1998; Kruuk 2004). This shows whether offspring of related mothers share similar trait values (whilst accounting for the additive genetic effects), and thereby assigns a portion of the variance to maternal genetic effects. Thus with maternal genetic (*V_Ma_*) and environmental (*V_Mn_*) variance, the total variance is:

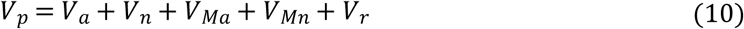

#### Variance estimates for swimming speed

Using a model with only additive genetic and residual variance, both components were overestimated *V_a_* = 2.067 (SE=0.22, p<0.001), *V_r_* = 2.082 (SE=0.13, Fig 5a) and as a consequence a large heritability was found *h*^2^ = 0.498 (SE=0.038).

**Fig 5:**
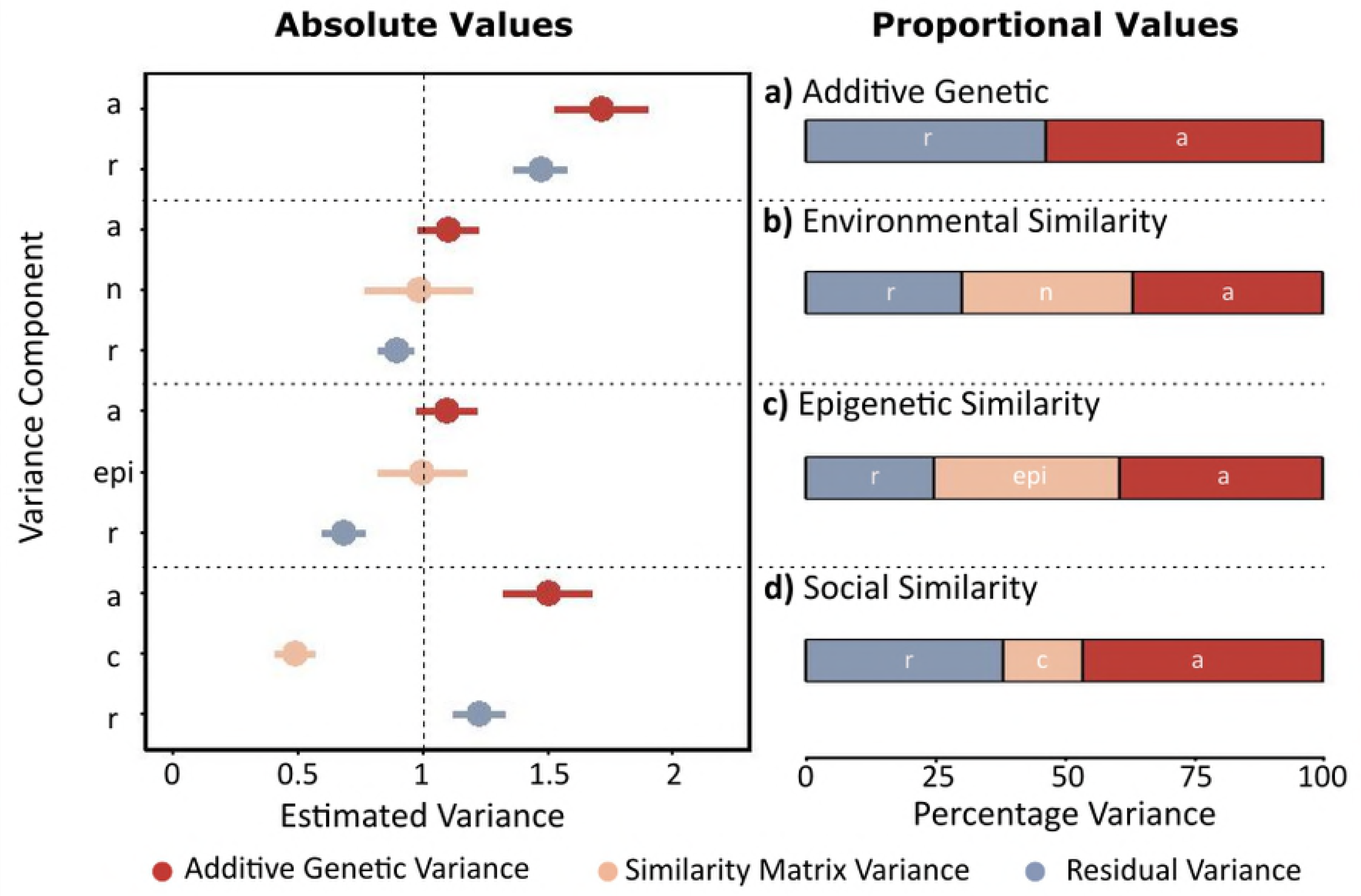
Variance components of trait 3. This trait was simulated to contain additive genetic, maternal genetic, environmental, maternal environmental, and residual variance, each with mean=0 and variance=1. For each of the models, the pair of plots show the absolute values (left plot) and the proportion (right plot) of total trait variance explained. a) Model containing only additive genetic (*a*) and residual variance (*r*). b) Model including additive genetic (*a*), environmental (*n*), and residual variance (*r*), and maternal variance estimated by including maternal identity as a random effect (*M*). c) Model including additive genetic (*a*), maternal genetic (*Ma*), environmental (*n*), maternal environmental (*Mn*), and residual variance (*r*)(the true model).

A model that included direct environmental variance, and maternal variance (using maternal identity as a random effect) improved estimation (Fig 5b), but *V_a_* remained overestimated (*V_a_* = 1.459, SE=0.22, p<0.001). A large maternal effect was found (*V_M_*= 1.302, SE=0.15, p<0.001), although it underestimated the total maternal variance, which is expected to be ~2. Direct environmental variance was estimated correctly (*V_n_* = 1.094, SE=0.27, p<0.001). Overestimation of components in this model arose because maternal variance was simulated to be both environmentally and genetically derived, and so the complete model must estimate both maternal genetic variance, and maternal environmental variance.

The full animal model included additive genetic, maternal genetic, direct environmental, and maternal environmental variance (equation 10). This model revealed a significant maternal genetic variance *V_Ma_* = 0.792 (SE=0.14, p<0.001), although this was slightly underestimated, and maternal environmental variance *V_Mn_* = 1.345 (SE=0.41, p<0.001), which was slightly overestimated. Other components were well estimated, *V_a_* = 1.119 (SE=0.19, p<0.001), *V_n_* = 1.118 (SE=0.26, p<0.001), *V_r_* = 0.922 (SE=0.11), with heritability *h*^2^ = 0.211 (SE=0.038).

This model follows the simulation of this trait. Whilst the direct genetic and environmental effects were well estimated, the maternal variance components were not as well estimated (Fig 5b). The maternal environmental variance was overestimated, whereas the maternal genetic variance was underestimated. This is due to the difficulty in distinguishing between the two maternal effects. These are challenging to disentangle, because related mothers are also likely to have similar environments. Whilst this is difficult to address with wild populations, increased data sizes (see Data Requirements) may aid in estimation. Experiments may also be of use, such as experimentally increasing dispersal between environments.

### Epiallele similarity matrix

Using the first two traits (tail-fin colour and body size), we now show how variance in a trait could be partitioned using matrices derived from other possible sources of variance. We do not consider swimming speed (trait 3) from this point onward because of the difficulties estimating maternal environmental effects demonstrated above. Epigenetic information was converted into an epigenetic similarity matrix (**S_epi_,** Fig 2c). As with the environmental information, the value for each island was first scaled so that it had a mean of zero and variance of one. From this scaled epigenetic information, the Euclidean distances between all pairs of individuals were calculated (as in Johannes *et al.* 2009), and then scaled to be between 0 and 1. Thus, the elements of the matrix are:

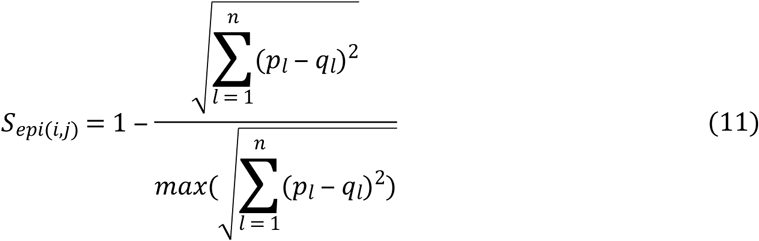

where *p_l_* and *q_l_* are the vectors of scaled epigenetic measures for individuals *i* and *j*, akin to the elements of the **S_n_** matrix (equation 6). We expect that this matrix would capture similar information to the environmental similarity matrix above - individuals that experience similar environments will be more similar in their epigenome than those in dissimilar environments. To distinguish between genetic and epigenetic variances, there need to be epigenetic differences between relatives. If all epigenetic changes are obligatory (i.e. genotype dependent; Richards 2006), epigenetic variance is simply nested within the genetic variance.

Using the epigenetic similarity matrix, we can partition phenotypic variance into genetic and epigenetic variance parameters (*V_epi_*):

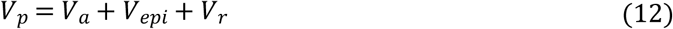

For traits including maternal epigenetic variance (*V_mepi_*):

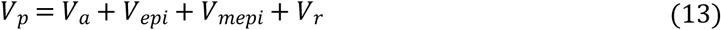

Thus, these models estimate whether individuals that have similar epigenetic profiles have similar trait values. For tail-fin colour, this model gave variance estimates of *V_a_*= 1.097 (SE=0.12, p<0.001), *V_epi_* = 0.998 (SE=0.18. p<0.001), and *V_r_* = 0.684 (SE=0.09), with a heritability of *h*^2^= 0.395. Thus, the epigenetic similarity matrix accurately recapitulates the environmental effects on the trait (Fig 3c).

For body size, this model gives variance estimates of *V_a_*= 1.091 (SE=0.16, p<0.001), *V_epi_* = 1.085 (SE=0.2, p<0.001), *V_Mepi_* = 0.699 (SE=0.12, p<0.001), and *V_r_* = 0.614 (SE=0.1; Fig 4e). Heritability is *h*^2^ = 0.313 (SE=0.042). Thus, there is some underestimation of the maternal variance using the epigenetic similarity matrix, which is likely to be due to the fact that the trait was not simulated directly on the epigenetic effects.

Currently, it is be challenging to obtain this sort of information for wild populations. However, we hope that the feasibility of obtaining this data will improve over time, in the same way that genotyping has. Therefore, the technique we present here will contribute to a growing area of research. Using matrices of epigenetic similarities could also be useful in laboratory or experimental situations. In cases where there is no genetic variation between individuals (e.g. using clones), data requirements will be lower, as *V_a_* = 0 by definition.

It should also be noted that in this simulation, using these alternative matrices technically produces type 1 errors (false positives), as tail-fin colour and body size were not simulated to be directly affected by either epigenetic or social effects. However, these models work as a proof of concept.

### Social network matrix

To include social network information in a model, we used non-weighted connections between individuals to generate the **S_c_** matrix of geodesic distances between individuals (i.e. the shortest path *g_i_*,*_j_* between individuals *i* and *j*; Fig 2d). Matrix diagonals must be one, so we adjusted the geodesic distances to have a path length of one to themselves, and added one to all other distances. We then took the reciprocal of each element, so that closely connected pairs had values close to one. Thus the elements of the **S_c_** matrix are:

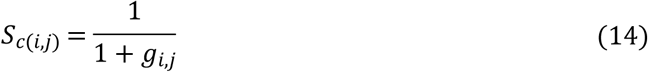

A matrix that describes individuals’ proximity or connectedness in a social network can therefore be passed to the model to account for this:

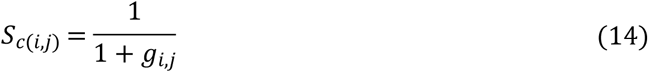

where *V_c_* is variance in the direct effect of the social network similarity on trait values. This model therefore estimates how much of the trait can be attributed to proximity of individuals in their social network.

Parental social network variance can also be estimated using the **S_c_** matrix:

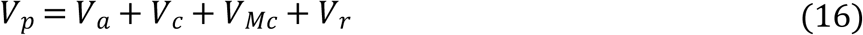

where *VM_c_* is the variance associated with the maternal effect due to her social interactions.

For tail-fin colour, we found a small but significant amount of variance due to social connections (*V_c_*= 0.491, SE=0.08, p<0.001). This was an underestimate of the variance due to nongenetic causes, and as a result the additive genetic variance *V_a_* = 1.501 (SE=0.18, p<0.001), residual variance *V_r_*= 1.226 (SE=0.11), and heritability *h*^2^ = 0.467 (SE=0.044) were overestimated (Fig 3d).

For body size, we found *V_c_*= 0.265 (SE=0.09, p<0.001) and *VM_c_* = 0.622 (SE=0.13, p<0.001), showing that there was significant variance estimated by the social similarity matrix. Again, these values were underestimated relative to the true variance, and so the additive genetic *V_a_* = 1.213 (SE=0.2, p<0.001) and residual variance *V_r_* = 1.492 (SE=0.13) were larger than one (Fig 4f).

To partition variance attributable to the social network, there must be sufficient contrast between the genetic and social similarity matrices. For example, in a social network in which connections only occur between parents and offspring, it will be difficult to disentangle social and additive genetic variance. If there are both full- and half-siblings within the data, and differences in the transmission of the trait from mothers and fathers, the social effect can be estimated as part of the maternal or paternal effects. More generally, social information can be incorporated if related individuals are present in different social groups, and differ in their connections in the network.

The matrix of social connectedness between individuals estimates some, but not all, of the environmental variance in the data. This is due to two main reasons. Firstly, the trait was simulated over the continuous environmental measures, not the social network, which was simulated using the spatial information for individuals. It is therefore unsurprising that it fails to capture all of the variance. Secondly, connectivity decreases with generational distance but environmental (and epigenetic) similarity does not. A trait that was truly inherited through a social network would be likely to show a more precise estimate of social variance.

## Discussion

In this paper, we outlined how to simultaneously decompose genetic and nongenetic (environmental, epigenetic, and social) sources of similarity between individuals in an animal model. We also provided a supplementary tutorial, using existing software, to encourage the use of these methods. This should facilitate wider adoption of these methods by researchers investigating phenotypic variation in wild or laboratory populations.

With biologically plausible simulated data, we estimated significant genetic, environmental, epigenetic, and social variation in all three traits. As has been shown previously, *V_a_* estimates of these traits changed significantly when variance due to nongenetic effects was estimated, leading to more accurate estimates of additive genetic variance and heritability (Kruuk & Hadfield 2007; Noble *et al.* 2014). For studies of wild populations, the actual effects of including nongenetic similarity matrices on estimates of variance will depend on the trait in question - as seen in the various effects of including spatial similarity in Stopher *et al.* (2012), Regan *et al.* (2017), and Germain *et al.* (2016).

The relative effect of including of nongenetic similarity on *V_a_*, *V_r_*, and *h*^2^, will depend on whether the causes of similarity increase the resemblance between relatives or non-relatives. Nongenetic resemblance amongst relatives artificially inflates a trait’s estimated additive genetic variance when not accounted for. Conversely, increased resemblance between unrelated individuals will affect estimates of the residual variance, but heritability should be unbiased. In the merpeople simulation, relatives were similar in their continuous environmental measures, so that genetic variance was inflated by environmental effects when they were not accounted for.

For body size and swim speed, maternal effects were also included in the models, showing that it is important to account for maternal traits when they affect the trait of interest. For traits where maternal effects don’t exist, or have no effect on the estimation of additive genetic variance, the inclusion of maternal effect variance is unlikely to change estimates of additive genetic variance. Maternal environmental and genetic variance proved challenging to partition completely, but neglecting maternal environmental variance might cause over-estimation of maternal genetic, as well as additive genetic, variance.

### Challenges with nongenetic matrices

Inclusion of such matrices with real-world data should be hypothesis driven, such that researchers have reason to expect that nongenetic similarity may account for some portion of phenotypic variance. These matrices are non-sparse, and the models run more slowly and require more computer memory than an animal model containing only the additive genetic relatedness matrix.

Multiple nongenetic parameters themselves can have genetic influences (e.g. habitat selection, dispersal, social behaviours). For example, if there is a genetic basis to social behaviour (Fowler *et al.* 2009), the social network and genetic variance could interact in forming the trait of interest. Including a behavioural trait or network metric (e.g. dominance) as a second response variable in a bivariate model could help account for this interaction. In addition, there may be interactions amongst sources of (nongenetic) variance. For example, Champagne & Meaney (2006) showed that a genetically determined maternal effect (maternal care) created a maternal environment that led to the methylation of rat pup DNA. Therefore, the maternal genetic and direct epigenetic effects are covarying and confounded, and the estimation of the epigenetic effect may depend on whether a maternal genetic effect is included, and vice versa (*sensu* causally covarying traits, Morrissey 2014; Hadfield &Thomson 2017).

The additive genetic variance and heritability of traits, can change over ontogeny (e.g. Wilson *et al.* 2005b) or senescence (e.g. Pujol *et al.* 2014), but the additive genetic relatedness between individuals does not generally change over time. This is not necessarily the case for nongenetic factors. These may change over a lifetime in both the variance explained (as with additive genetic variance), and changes in the similarity matrix itself. The environment (Charmantier *et al.* 2008, Charmantier & Gienapp (2014)), epigenetic marks (Polanowski *et al.* 2014) and social networks (Wey & Blumstein 2010) can all change over time, potentially altering the similarity between individuals. Furthermore nongenetic similarity during development may explain more variance than nongenetic similarity measured at the same time as the phenotype (or vice versa). For example, Pigeon *et al.* (2017) showed the importance of early-rather than later-life environment on traits, and Regan *et al.* (2017) used the home range overlap of mothers to investigate offspring early-life traits. The relative importance of similarity at different times could be investigated using multiple nongenetic similarity measures to test age-specific effects.

Carrying out these analyses may also be made more challenging by the presence of missing data for the nongenetic components. Stopher *et al.* (2012) included individuals with missing data in the model, allowing a similarity of 0 to all other individuals in the spatial overlap matrix (indicating no overlap). Our data has missing data (environmental, epigenetic, and social) for the parents of the first generation of individuals, and so these parents are not included in the nongenetic similarity matrices (including the parents, but with similarities of 0 to all individuals does not qualitatively change the variance components in the model). In cases where some (but not all) environmental (or epigenetic) data is missing for an individual, it may be possible to impute the values based on the mean environmental value, either at a dataset-wide or smaller spatial scale. Finally, caution must be taken about the potential effect of errors in nongenetic data. The effect of errors will be dependent on their frequency and whether there are random or systematic errors. Random errors should have relatively little effect, but systematic errors and those that correlate with an underlying trait may cause biased estimates (akin to non-random errors in paternity; Firth *et al.* 2015).

### Environmental similarity

Environmentally induced resemblance between relatives will depend upon the heterogeneity of the environment, and the rate and distance of dispersal within and between environment types. For example, in cases where dispersal away from the mother is low relative to the rate of environmental change, full- and maternal half-siblings are likely to share both genes and their environment, increasing their resemblance. In the extreme case, this may lead to the two variance components being completely confounded. Cross-fostering (Merilä & Sheldon 2001; Danchin *et al.* 2013; Winney *et al.* 2015), transplantation (Kawecki & Ebert 2004; Fang *et al.* 2006), or laboratory experiments that fix certain sources of variance can help separate these effects from genetic variance.

Environmental measures can have non-linear and interacting effects on phenotypes. Fitting separate environmental measures as fixed effects makes assumptions about the linearity or interactions of such effects. These assumptions are not made with the environmental similarity matrix, though it does assume that its component environmental measures have equal weight. Thus the choice of how to include environmental measures in an animal model may depend on the questions researchers want to address. For example, tail-fin colour in merpeople could be strongly affected by salinity and less strongly affected by other environmental measures, which could potentially water down the strength of the covariance between the environment and the phenotype. If we were interested specifically in how salinity affects tail-fin colour we could choose to fit salinity as a fixed effect, rather than incorporating it into the matrix with the other nine environmental measures. For factorial environmental measures, these could be fitted as an additional random effect. Conversely, if we wished to estimate environmental variance dependent on all the measures, then we would retain it within the matrix.

### Epigenetic similarity

The simulation that we show here is an oversimplified view of how epigenetics could be measured, but is used here to illustrate how such effects could be accounted for. We use a simulation of methylation at CpG islands (Gardiner-Garden & Frommer 1987; Jones & Takai 2001), but there are many more forms and states of epigenetic variation that could be measured, such as methylation states at individual loci, histone modification (acetylation, methylation, phosphorylation, and ubiquitination), and RNAis (Bossdorf *et al.* 2008; Kovalchuk 2013). Additionally, only one type of epigenetic measure is included - it assumes all cells are the same in epigenetic marks, or that we have only measured epigenetic marks in cells relevant to the trait. Alternatively, data used from Epigenome-Wide Association Studies (EWAS) could be used, wherein the matrix could contain information in the similarity between individuals in their percentage methylation at multiple loci across multiple cell types. Furthermore, the magnitude of the epigenetic effect in our simulation could depend upon the rate of resetting or reconstruction between generations. Thus, establishing the rate of reset or reconstruction is important in understanding how resetting affects quantitative genetic estimates and evolutionary processes (Tal *et al.* 2010; Kovalchuk 2013; Kronholm & Collins 2016).

Additionally, the genomic architecture of epigenetic marks may be complex. For example, epigenetic marks could be in linkage disequilibrium with genetic variance (Taudt *et al.* 2016), particularly if the genotype of an individual determines the probability of certain epigenetic states (facilitated epigenetics; Richards 2006). This makes drawing apart genetic and epigenetic variance challenging, and makes studies that examine epigenetic variation in a fixed genetic background valuable (e.g Cubas *et al.* 1999; Latzel *et al.* 2012; Cortijo *et al.* 2014). Such studies could make use of the models outlined here, excluding relatedness data, to estimate variance due to different nongenetic parameters (e.g. direct *vs.* maternal epigenetic effects).

### Evolutionary implications

Nongenetic effects may directly affect evolutionary potential (Danchin *et al.* 2011), and may be involved in the missing response to selection of wild populations (Pujol *et al.* 2018). The response of heritable nongenetic variation to environmental change can increase the rate of both phenotypic and genetic responses (Bonduriansky *et al.* 2012), and may therefore affect population persistence. However, the concept of nongenetic inheritance remains controversial and poorly understood. Theoretical progress has been made on this topic (Bonduriansky & Day 2009; Day & Bonduriansky 2011; Townley & Ezard 2013), though empirical evidence is lacking in wild systems. The evolutionary importance of nongenetic inheritance relies on several prerequisites that need to be empirically evaluated. Firstly, nongenetic changes must be transmissible. Secondly, these changes must be (partially) stable over multiple transmission steps. Finally, nongenetic inheritance must affect evolutionary potential.

Multiple matrix models can provide novel insights for these problems: first, they allow us to demonstrate consistent nongenetic effects on the phenotype, and quantify the proportion of variance explained by these nongenetic effects. Second, where there is separate evidence for the stable transmission of the nongenetic effect (e.g. epigenetic inheritance), the model will show the relative effect of these nongenetic mechanisms compared to genetic mechanisms. For the traits simulated in this paper, a portion of the epigenome is maternally inherited, which in turn implies that a portion of *V_epi_* is inherited. This animal model containing epigenetics does not distinguish between the portion of variance that is due to inherited (directly or reconstructed) or non-inherited epialleles, but it does imply that nongenetic inheritance may be having a significant effect on trait variation. If heritable epialleles are known *a priori*, these could be included as a separate matrix, allowing distinction between heritable and non-heritable epigenetic variation. In addition, comparison of trait covariances between different types of relatives may reveal the rate of epigenetic inheritance and reset (Tal *et al.* 2010)
.

Furthermore, in a population where individuals show close affinity to parental environments, such as inheriting a territory (e.g. red squirrels; Berteaux & Boutin 2000), niche transmission may form a part of the estimate of environmental variance (*V_n_*, and potentially *V_Mn_*; Odling-Smee & Laland 2011). Niche transmission cannot be implied directly from the animal model containing the environmental similarity matrix, but complementary external evidence may be used to support such a hypotheses in some populations. Similarly, estimates of variance due to social network connectivity may be used in conjunction with other information to make conclusions about social inheritance or cultural transmission of traits. For example, if teaching is known to happen between mothers and offspring, or within peer groups, then some portion of *V_c_* is likely to represent the inheritance of the trait, rather than just that attributable to similarity.

### Conclusion

A major advantage of the multiple matrix approach is its flexibility in where and how it can be applied. Nongenetic similarity matrices are not restricted to those demonstrated in this tutorial, but could also include a range of other measures of similarity, for example gene expression (Kim *et al.* 2007), histone states (Taudt *et al.* 2016) or gut flora communities (Vries *et al.* 2001). Consequently, this approach has the potential to bring together disconnected fields, such as ecology and molecular biology, which would connect information on broad-scale variation in natural systems to mechanistic information gained from model systems (Richards *et al.* 2017).

The multiple matrix extension to the animal model has potential applications to both wild and laboratory data. Wider adoption of multiple-matrix models has substantial promise for improving our understanding of ecological and evolutionary processes. The complex interaction of genetic and nongenetic sources of variance with dispersion, the rate of loss or reset of nongenetic information, and population structure may help to explain some of the differences between populations in estimates of genetic variance (Teplitsky *et al.* 2014). Therefore, the magnitude of nongenetic effects on evolutionary potential has not been quantified, but has the potential to be substantial.

## Acknowledgements

We thank Laura Gervais, Delphine Gourcilleau, Mathieu Latutrie, Pascal Marrot, Mathilde Mousset, and Charlotte Regan for useful feedback. We are also grateful to Matthew Silk for aid with social network simulation and analysis. Additionally, we thank Alastair Wilson and Matthew Wolak for helpful reviews. This work is part of a project that received funding from the European Research Council (ERC) under the European Union’s horizon 2020 research and innovation program (grant agreement ERC-CoG-2015-681484-ANGI to BP). This work was also supported by the French “Agence Nationale de la Recherche” (ANR-13- JSV7-002 “CAPA” to BP). CET, ISW, OCS and BP were also supported by the ANR funded French Laboratory of Excellence projects “LABEX TULIP” and “LABEX CEBA” (ANR-10- LABX-41, ANR-10-LABX-25-01) and ANR funded Toulouse Initiative of Excellence “IDEX UNITI” (ANR11-IDEX-0002-02).

## Authors’ Contributions

All authors conceived the ideas and designed the methodology. CET simulated and analysed data. CET, ISW, OCS wrote the manuscript. BP provided feedback and editing.

## Data accessibility

All analyses are freely available as the supplements to this paper, as is the code used to create data. Data will be archived at Zenodo upon acceptance.

